# Blinded by motion: Electrophysiological decoupling of time dilation and confidence

**DOI:** 10.64898/2026.09.18.752748

**Authors:** Tutku Öztel, Martin Wiener

## Abstract

The duration of visual stimuli can be altered by stimulus features, such as size, brightness, and motion. Recently, we observed that motion-induced changes in perceived duration also shift confidence judgments, but do not track awareness of the illusion. How these two effects differ and are instantiated neurally are unknown. We investigated this question by replicating our previous finding, in which human subjects (n=27) performed a temporal bisection task where they were asked to classify the duration (1-3.5s) of a walking stickman figure with different velocities and provide confidence judgments while 64-channel electroencephalogram (EEG) activity was recorded. Behavioral findings replicated the previous time dilation effect where progressively faster walking speeds led to longer perceived durations. A similar shift was observed for confidence ratings, yet with lowest confidence for the medium walking speed, indicating a metacognitive inability to monitor dilation. Furthermore, common EEG signatures of time perception, such as the late positive component of timing (LPCt) and contingent negative variation (CNV) did not vary with walking speed. Instead, onset-locked responses over occipital electrodes covaried with walking speed and correlated with subject-level shifts in psychometric functions, but not with confidence; conversely, frontocentral responses covaried with confidence judgments, but not walking speed. These findings suggest that temporal illusions are engaged by sensory processes that fail to reach the metacognitive level and that confidence judgments, in turn, are driven by motor estimates read out from frontocentral regions.

## Main

Human time perception is subjected to distortions from stimulus psychophysical properties associated with the sensory environment that abides a “more A more B” perceptual pattern (Walsh, 2003; Bueti & Walsh, 2009; Winter et al, 2015). One robust time dilation effect is induced via stimulus velocity (e.g., Carrozzo & Lacquaniti, 2013; Karsilar, Kisa & Balci, 2019; Oztel & Balci, 2020; also see: Mioni, Zakay & Grondin, 2015). While humans are aware of their motor timing errors on a trial-by-trial basis (e.g., Akdogan & Balci, 2017; Oztel, Eskenazi & Balci, 2020; for a detailed review: Oztel & Balci, 2024), this metacognitive ability does not readily translate to dilated temporal representations with walking speed (Oztel & Balci, 2020). This raises the possibility that stimulus induced time dilation and metacognitive representations might be distinctly represented in the brain. The current study investigates the neural basis of metacognitive processing of time dilation induced by the velocity of animating stimuli.

Previous research on the neural basis of metacognition in perceptual decision making shows correlation between the subjective confidence ratings and electroencephalogram (EEG) signals. For example, post-error negativity (Pe) amplitude scales negatively with decision confidence (Boldt & Yeung, 2015). Similar results associated with different event-related potential (ERP) signals were reported in many other studies (e.g., frontocentral error related negativity (ERN) and Pe; Kirschner, Humann, Derrfuss, Danielmeier, & Ullsperger, 2021; see also: Ko, Zhou, Niessen, Stahl, Weiss, Hester,… & Feuerriegel, 2024; Stone, Mattingley, Bode, & Rangelov, 2024; Zakrzewski, Wisniewski, Iyer, & Simpson, 2019; Salti, Bar-Haim, & Lamy, 2012; Selimbeyoglu, Keskin-Ergen, & Demiralp, 2012; for a detailed review: Wessel, 2012; Fleming, 2024). Critically, the frontocentral ERN signals observed in these studies are thought to be sourced from the anterior cingulate cortex (ACC) to encode conflict monitoring processes (Botvinick et al., 2004), which has further support from more direct neuroimaging techniques with enhanced spatial resolution (like fMRI: Hebart et al., 2016; Morales et al., 2018; dorsal ACC and preSMA; Fleming et al., 2012; Heekeren et al., 2008; Ridderinkhof et al., 2004, dACC and SMA: Katayama et al., 2025). Taken together, these results highlight the critical involvement of ACC in metacognitive and conflict monitoring processes.

Similar to metacognitive processes, time perception has also been documented to have frontocentral associations, particularly with supplementary motor area (SMA; e.g., Macar et al., 2006; Coull et al., 2015; for a review: De Kock, Gladhill, Ali, Joiner & Wiener, 2021), such that the contingent negative variation (CNV) amplitude from SMA is correlated with evidence accumulation for the perceived duration (e.g., Macar, Vidal & Cassi, 1999; Ng et al., 2011; Baykan et al., 2023; Casini & Vidal, 2011) and other time-related processes including sensory expectation and motor preparation associated with time-related decision processes (e.g., van Rijn et al., 2011; Kononowicz et al., 2011; 2014; 2016). Another frontocentral ERP component associated with timing behavior is the late positive component of timing (LPCt), which is known to encode decision making processes and discrimination difficulty (Paul et al., 2003; 2011; Gontier et al., 2009). Together, these results suggest that time perception and metacognition have spatially close (i.e., frontocentral) but distinct neural correlates.

In the current study, we investigated the neural correlates of stimulus induced time dilation and potential metacognitive neural markers associated with them in a temporal bisection task. Participants were asked to classify the duration of a walking stickman figure in different velocities (slow, medium and fast) as being closer to a short or long reference duration and reported their confidence (as in Öztel & Balci, 2020). During the experiment, we recorded participants’ brain activity with EEG. We hypothesized that if the metacognitive inability associated with the stimulus induced time dilation effect is due to their distinct representations, these two cognitive processes should be represented in distinct regions in the brain. This potential distinction should reflect itself in differential relationship between ERPs and leftward shift in the psychometric functions associated with different stimulus velocities (as an index of time dilation) and confidence ratings (as an index of metacognitive monitoring, see: Fleming, 2012).

## Method

### Participants

28 neurologically and psychiatrically healthy students from George Mason University with normal or corrected-to-normal vision participated in this study. One participant did not complete the experiment due to motion nausea. As a result, the final analyses were conducted on the remaining 27 participants (*M*_age_ = 24.07, *SD*_age_ = 5.28, 15 females, 1 nonbinary, all right handed). All participants provided written consent to participate in the current study and were monetarily compensated. All protocols were approved by the Institutional Review Board of George Mason University.

### Stimuli and Apparatus

All the stimuli used in the current study were the same as in Oztel and Balci (2020, see also: Karsilar, Kisa, Balci, 2019). The stimuli set consisted of forward walking stickman figure videos with three different walking speeds (25 (slow), 50 (medium), 100 (fast) frames per second; at 1, 1.5 and 3 Hertz frequencies, respectively) and six different probe durations (1, 1.5, 2, 2.5, 3, 3.5 seconds). The stimulus size was set to 0.5 x 0.5 in height unit compared to the screen size. As a result, the stimuli set consisted of 3 (speed) x 6 (duration) = 18 different walking stickman figure videos in total. All responses were made with a standard computer keyboard. The experiment was coded in PsychoPy Builder mode (version: 2024.2.4, Peirce et al., 2019). All the stimulus presentations and response recordings were performed with PsychoPy. Participants sat in front of a 32″ LCD Monitor (Cambridge Research Systems Display++) running at 120 Hz refresh rate with 1920 × 1080 resolution at a distance of ∼70 cm.

### Procedure

The behavioral task of the experiment was a temporal bisection task. Participants were instructed to respond as fast and as accurately as possible.

#### Training Phase

Participants were first presented with both short (1 second) and long (3.5 seconds) anchor durations in a randomized order with a square noise patch (0.2 x 0.2 in height unit compared to screen size). Then, they were randomly presented with either of the durations and asked to classify the presented duration as being the short or long reference duration. After each of their classifications, participants received feedback for their classification accuracy. The training phase lasted 10 trials.

#### Testing Phase

Participants were presented with both the anchor durations and probe durations with animated walking stickman figures with different walking speed. Participants classified the duration of the animated stimuli as being closer to the short or long anchor duration by pressing “S” and “L” for “short” and “long” responses, respectively (self paced). Each classification was followed by a 500 millisecond central fixation cross. After each of their responses, participants were asked to rate their level of confidence regarding the accuracy of their classifications (“1”-low confidence - “3”-high confidence, self paced). Each trial ended with 1-1.5 second inter-trial interval (ITI) randomly chosen from a uniform distribution. There were three blocks separated with one minute of break. Each block consisted of 216 trials. As a result, the experiment consisted of 216 x 3 = 648 trials in total and took around 70 minutes to complete. Figure 1A illustrates an example trial in the experiment.

**Figure 1.**
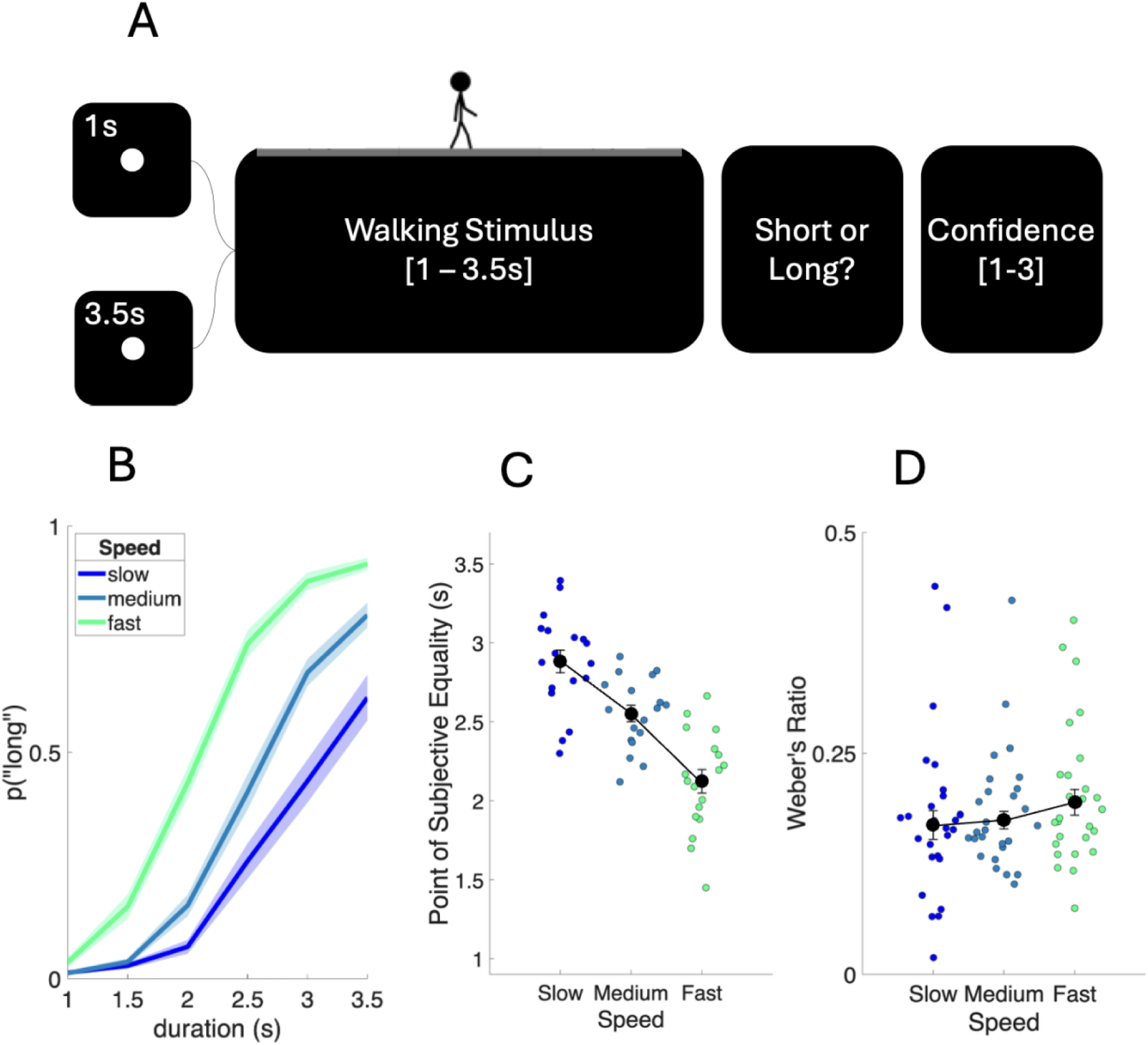
The experimental procedure and the behavioral results for the time dilation as a function of walking speed. (A) Schematic representation of the experiment. Participants were presented with a walking stickman figure with different speeds and durations and asked to classify its duration as “short” or “long”. After each response, they reported their confidence. (B) p(“long”) responses across stimulus durations and walking speeds (C) Point of subjective equality (PSE) values across slow, medium and fast walking speeds. (D) Weber’s ratio (WR) values across slow, medium and fast walking speeds. In both panels, scatterplots indicate individual observations associated with each participant in each walking speed condition. Error bars and shaded areas represent standard error (SE)

#### EEG recordings

EEG recordings were performed through a 64-channel actiCAP slim active electrode montage (international 10-20 system) with an actiCHamp amplifier (Brain Products GmbH, Germany). Throughout the recordings, FCz was used as an online reference. Prior to recordings, impedances associated with the electrodes were below 25 kΩ. Data processing was done with the EEGLAB toolbox for Matlab and epoched at the stimulus onset (i.e., onset locked). Prior to Infomax Independent Components Analysis (ICA), data were re-referenced to the offline reference electrodes (i.e., mastoid channels which are TP9 (back of left ear) and TP10 (back of right ear)). Data were then downsampled to 500 Hz. Data were low-pass filtered at 50 Hz and high pass filtered at 0.1 Hz. All noisy channels that were above 5 SD of the mean kurtosis for a joint probability distribution were removed. Eye blinks and other artifacts were removed from the epoched EEG data with ICA with runica via the built-in function for EEGLAB (Bell, Jung & Sejnowski, 1995; Makeig et al., 1996).

Before the behavioral experiment, we recorded participant’s baseline brain activity for five minutes during which they were instructed to look at a central fixation cross. Participants were not presented with any other stimulus or asked to make any responses during this period.

## Analytical Approach

### 1 Psychometric function fitting

We fitted standard Weibull Cumulative Distribution Function (CDF) to the probability of “long” responses separately for each walking speed and probe durations (i.e., *p*(“long”|duration, walking speed)). We then calculated the point of subjective equality (PSE) as the median of the psychometric function with the following equation, which corresponds to the duration that the participant perceives as long and short equally likely:

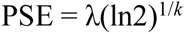

where λ indicates scale (i.e., location in the x axis) and *k* indicates shape (i.e., slope/steepness) parameter. We filtered out all data where PSE was smaller than 1 (i.e., the minimum stimulus duration) and larger than 3.5 seconds (i.e., the maximum stimulus duration) before the data analyses where PSE was included as either predicted or outcome variable.

We also calculated the Weber’s Ratio (WR) as the difference limen normalized with the PSE where

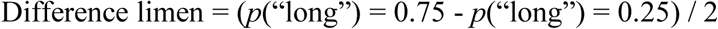

In the formula, the *p*(“long”) = 0.75 reads as “the duration that is classified as “long” for the 75% of the times” (= λ(-ln0.75)^1/*k*^). As a result, the WR is depicted as;

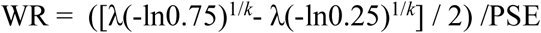

Accordingly, the lower values of WR indicates higher classification precision. Prior to data analysis, we discarded all WR values that were larger than 0.5 for all models where WR was used as a predictor or outcome variable.

### 2 Confidence ratings and Reaction Times (RTs)

#### Intersection points

We calculated the confidence intersection points in a way that indexes the equal confidence time point. For this, we ran an ordinary least squares (OLS) regression model using confidence as predicted and probe durations as predictor variables separately for short and long responses and three walking speeds. For each walking speed, we then calculated the intersection points of the two regression slopes associated with the short and long responses. Here, the intersection points refers to the duration which was rated with equal confidence. With these characteristics, the intersection points resemble the PSE (for a similar indexing, see Oztel & Balci, 2021). We calculated the RT intersection points in the same way as we did for confidence intersection points (i.e., the duration which was classified as “short” or “long” equally fast). We then investigated the linear relationship between the PSE and the confidence/RT intersection points. Accordingly, if the participants could keep track of their stimulus-induced temporal biases, the confidence and RT intersection points should not reliably increase with PSE (as in the case of Oztel & Balci, 2020). We discarded all data where confidence intersection was lower than 1 and higher than 3 (true confidence range) and RT intersection was lower than 0 and larger than 5, separately for each analyses where confidence and RT intersections were used as a predictor or outcome variable.

While investigating the relationship between the behavioral parameters and ERP amplitudes, we only used the ERP amplitudes from a time window that yielded a significant difference across the independent variables of interest for the given analysis (i.e., walking speed or stimulus duration) separately for onset locked epochs.

### Linear mixed effects model parameters

We investigated all linear relationships that included repeated observations with linear mixed effect models. For all linear mixed effects models, we used Satterthwaite as df approximation method and restricted maximum likelihood (REML) as parameter estimation method.

## Results

### Behavioral Results

#### 1 Leftward Shift in Psychometric Function as a function of walking speed

We investigated the leftward shift (i.e., PSE) and change in slope (i.e., WR) across in the psychometric functions different walking speeds with the following linear mixed effects model:

Model 1. PSE (or WR) ∼ walking speed + 1|participant

where PSE (or WR) was the outcome variable, walking speed was the categorical predicting factor with three levels (i.e., slow, medium and fast) and 1|participant indicated random intercept across participants. Linear mixed effects model revealed a significant main effect of walking speed (*F*(2,44.5) = 63.1, *p* < 0.001). Post hoc comparisons yielded a linear scaling of PSE as a function of walking speed (*M*_slow-medium_ = 0.201, *SE* = 0.0788, *p*_tukey_ = 0.037; *M*_slow-fast_ = 0.80, *SE* = 0.0788, *p*_tukey_ < 0.001; *M*_medium-fast_ = 0.60, *SE* = 0.0686, *p*_tukey_ < 0.001), indicating a significant leftward shift of the psychometric functions with walking speed (Figure 1B & 1C). On the other hand, WR values were comparable across different walking speeds (all *p*s > 0.05, Figure 1D).

#### 2 RT and confidence ratings as a function of walking speed

We further investigated how mean reaction times (RT) across different walking speeds and confidence ratings change as a function of walking speed with the following linear mixed effects model:

Model 2. mean RT ∼ walking speed + 1|participant

where mean RT was included as an outcome variable, walking speed was included as fixed effects and participants were included as random intercept in the model (i.e., “∼” reads as “predicted from” and “1|participant” reads as “random intercept across participants”). Linear mixed effects model revealed a significant difference across medium and fast (*M*_medium -fast_ = 0.0766, *SE* = 0.022, 95% CI = [0.033, 0.12], *p* = 0.001) and slow and fast speeds (*M*_slow - fast_ = 0.061, *SE* = 0.022, 95% CI = [0.0166, 0.105], *p* = 0.008), indicating a faster RTs for “fast” walking speed compared to medium and slow walking speeds. On the other hand, RTs were comparable for slow and medium speeds (*p* = 0.477). Figure 2A illustrates the mean RTs across different walking speeds.

**Figure 2.**
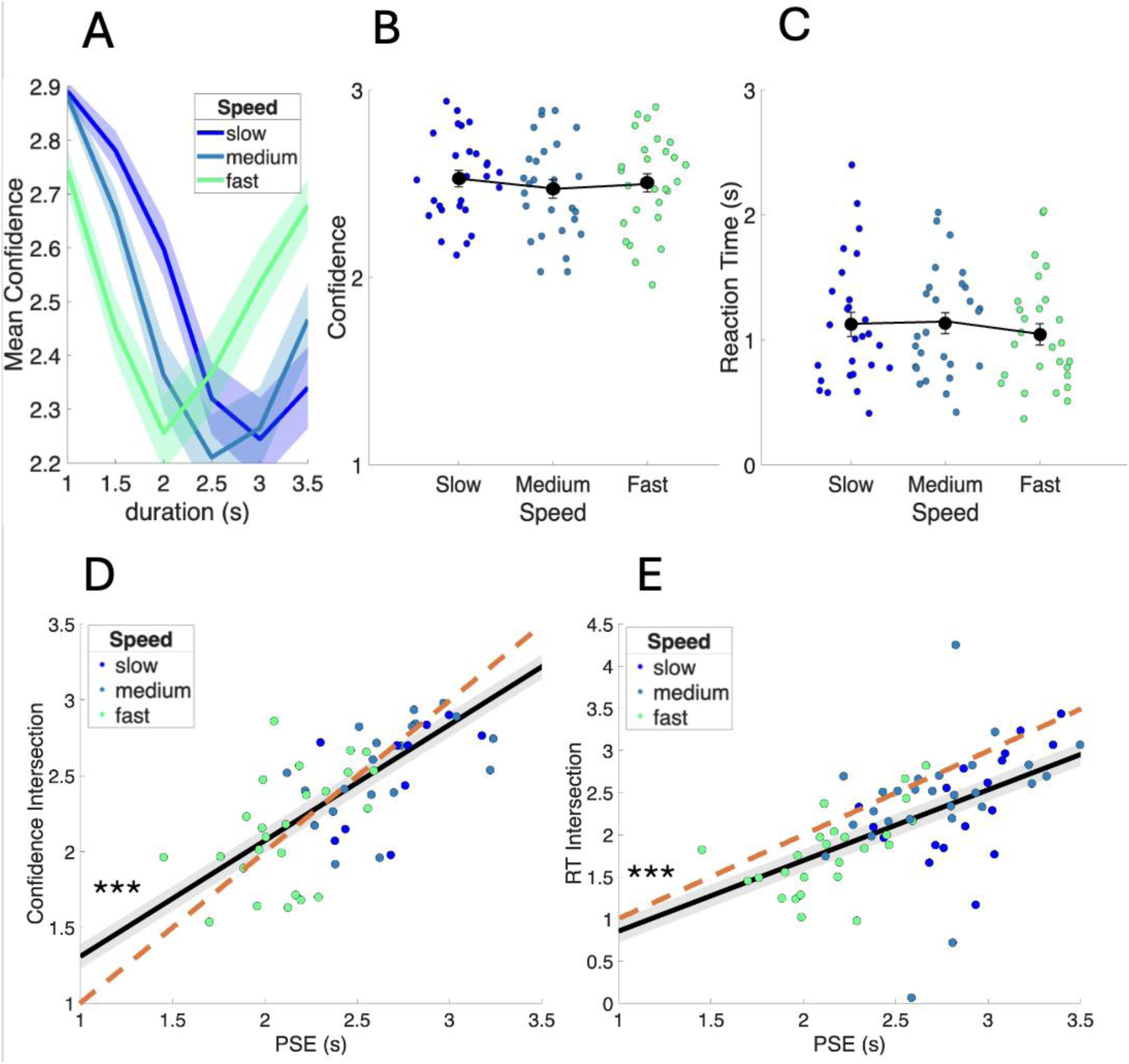
Behavioral results for confidence and reaction time (RT) across walking speeds and their correlation with shifts in psychometric function (PSE). (A) Mean confidence estimates across stimulus durations and walking speed (B) mean confidence ratings across three walking speeds (C) Mean RT across three walking speeds. Scatterplots indicate individual data points associated with each participant for each condition and data depicted in white circles indicate mean values. (D) The linear relationship between PSE and confidence and (E) RT intersection points. Scatterplots indicate individual data points for each walking speed condition. Error bars and shaded gray areas depict the standard error (SE). Dashed orange lines represent the y = x relationship. Asterisk indicate significance level (*** = *p* < 0.001).

We performed a similar comparison for confidence ratings, where we included mean confidence ratings across different walking speeds as an outcome variable and kept the remaining model parameters the same as in Model 1:

Model 3. Mean confidence ratings ∼ walking speed + 1|participant

Model 3 revealed a significantly larger confidence rating for slow, compared to medium walking speed (*M*_slow - medium_ = 0.053, *SE* = 0.019, 95% CI = [0.015, 0.092], *p* < 0.001) and for fast, compared to medium speed (*M*_medium - fast_ = −0.0314, *SE* = 0.019, 95% CI = [−0.070, −0.007], *p* = 0.005). On the other hand, confidence ratings were overall higher in slow, compared to fast speed (*M*_slow - fast_ = 0.022, *SE* = 0.011, 95% CI = [3.40e-4,0.0441], *p* = 0.047). These results reveal a nonlinear change in confidence ratings as a function of motion speed, which points to lowest confidence ratings for the medium walking speed. Figure 2B illustrates the overall change in confidence ratings as a function of walking speed.

#### 3 Confidence and RT Intersections as a function of PSE

We further investigated the relationship between the confidence and RT intersection points as a proxy of confidence equivalence for PSE (for a similar approach, see: Oztel & Balci, 2020) with the model depicted below:

Model 4. Confidence (or RT) intersection ∼ PSE + 1|participant

where confidence intersection points were an outcome variable, PSE was a fixed effect and participants were defined as random intercepts. Linear mixed effects model revealed a significant positive relationship between confidence intersection points and PSE (ß = 0.76, *SE* = 0.081, 95% CI = [0.60, 0.927], *p* < 0.001). A similar positive relationship was observed between PSE and RT intersection points (ß = 0.84, *SE* = 0.135, 95% CI = [0.57 1.108], *p* < 0.001). These results point out that participants cannot keep track of the shift in their PSE, which is in line with previous findings (Oztel & Balci, 2021). Figure 2D and 2E illustrates the relationship between PSE and confidence intersection and RT intersection, respectively.

### EEG Results

#### ERP amplitudes across stimulus durations

We investigated how average onset locked ERP amplitudes separately for occipital (O1, Oz, O2,PO7, PO3, POz, PO4, PO8) and frontocentral (FC1, Cz, FC2, F1, C1, C2, F2, Fz, FCz) electrodes changed as a function of stimulus durations. For occipital electrodes, onset locked ERP amplitudes peaked around the stimulus offset (monte carlo cluster corrected *p* < 0.05), demonstrating that the activity at occipital electrodes encode the physical onset and offset of stimulus durations (Figure 3A & B). On the other hand, the width of the ERP signal from frontocentral electrodes captures the subjective stimulus duration (monte carlo cluster corrected *p* < 0.05), suggesting that the activity at frontocentral electrodes encode the subjective readings of temporal information to predict stimulus duration (Figure 3D & E).

**Figure 3.**
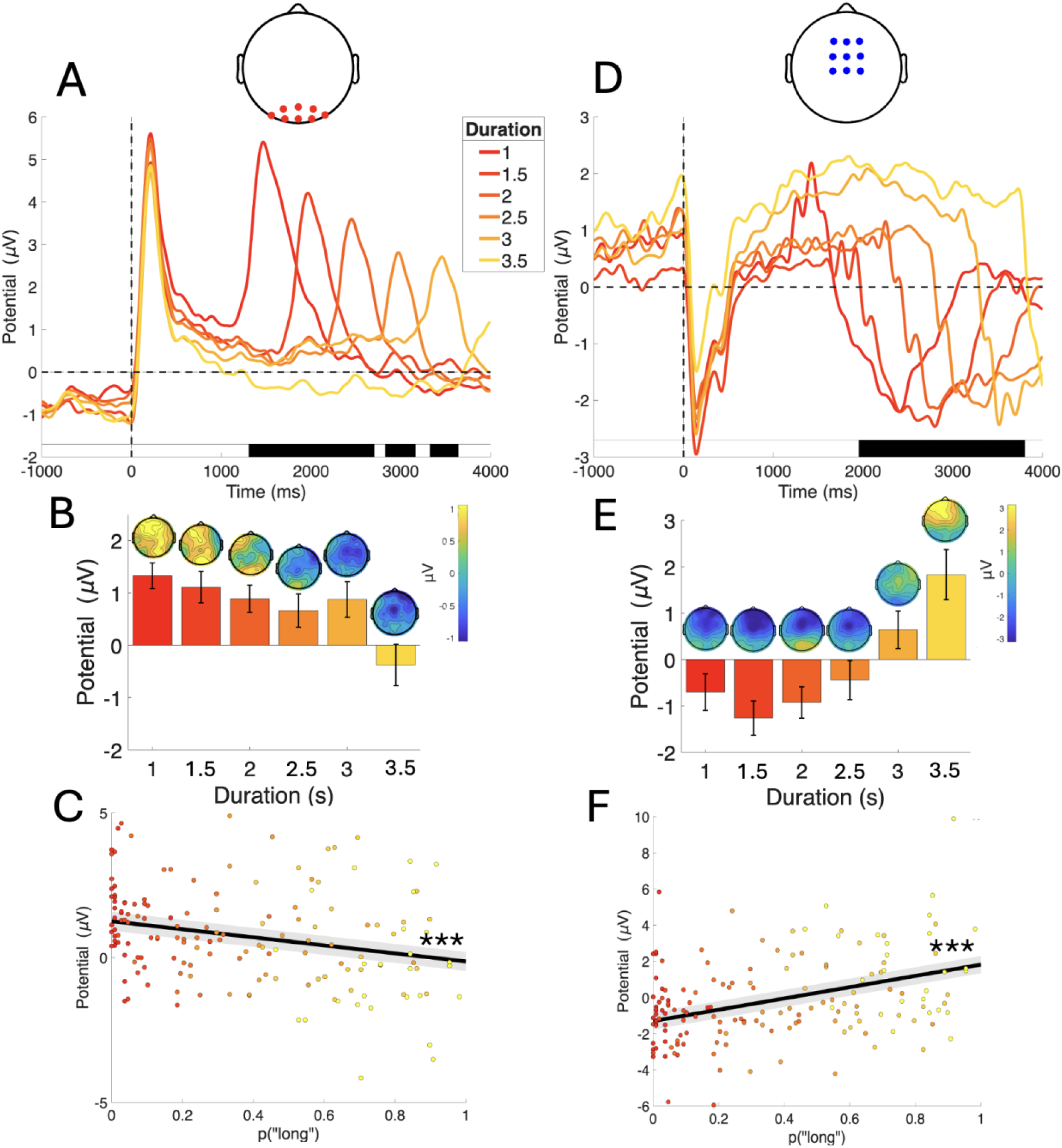
Duration splitted ERP signals for occipital and frontocentral regions. (A & B) The onset locked occipital and (D & E) frontocentral ERP amplitudes as a function of trial time for different stimulus durations (1-3.5 seconds, vertical dashed line indicates stimulus onset). The black bands indicate the time points where the amplitudes are statistically significant (monte carlo cluster corrected *p* < 0.05). Topographies illustrate scalp distribution of EEG activity at the stimulus duration. (C) The linear relationship between *p*(“long”) and onset locked occipital and (F) frontocentral ERPs. Scatterplots depict individual data across different stimulus durations. Shaded gray areas depict standard error (SE). Asterisk indicate significance level (*** = *p* < 0.001).

To validate this interpretation, we investigated the linear relationship between probability of “long” responses (i.e., *p*(“long”) associated with each stimulus duration:

Model 5. ERP ∼ *p*(long) + 1|participant

Linear mixed effects model revealed a significant positive relationship between mean frontocentral ERPs for each duration and the probability of “long” responses (ß_frontocentral_ = 3.12, *SE* = 0.48, 95% CI = [2.18 4.063], *p* < 0.001, Figure 3C), demonstrating that the ERP amplitudes for each stimulus durations increased as a function of “long” response frequencies. The opposite effect was observed for occipital ERPs, where ERP amplitudes associated with different stimulus durations decreased as a function of “long” response frequencies (ß_occipital_ = −1.39, *SE* = 0.33, 95% CI = [−2.04 −0.74], *p* < 0.001, Figure 3F).

#### ERP amplitudes across walking speeds

We investigated whether the ERP amplitudes from the occipital and frontocentral electrodes directly scaled with walking speed. Occipital ERP amplitudes indeed showed a direct scaling with walking speed, such that the largest amplitude was observed for fast walking speed whereas the smallest ERP amplitude was observed for the slow walking speed (monte carlo cluster corrected *p* < 0.05, Figure 4A). On the other hand, while the frontocentral ERP amplitudes were significantly different from each other for different walking speed (monte carlo cluster corrected *p* < 0.05), they did not directly scale with the walking speed, such that the largest ERP amplitude was observed for the medium walking speed (Figure 4D).

**Figure 4.**
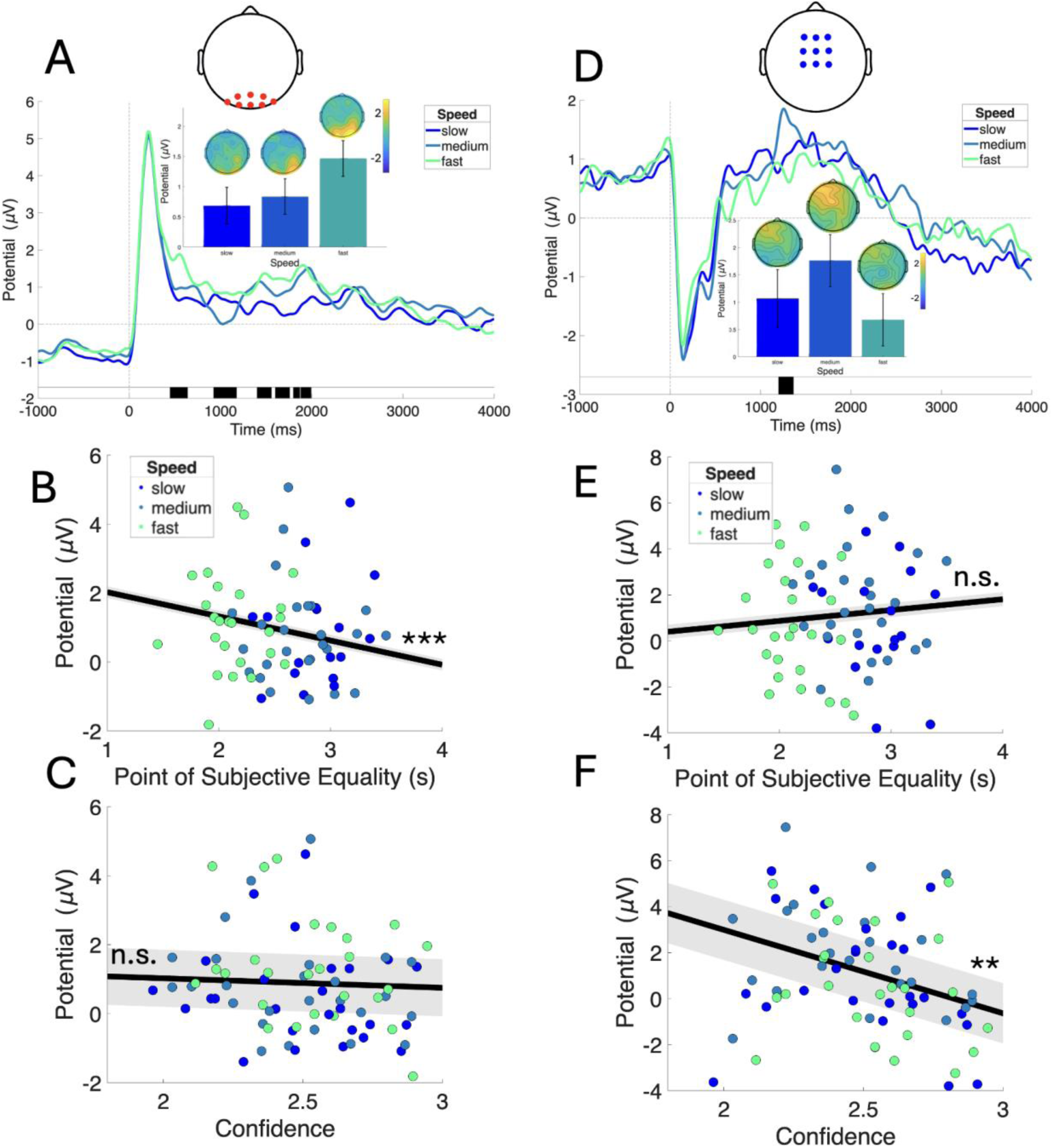
Speed splitted ERPs for occipital and frontocentral regions. (A) The onset locked occipital and (D) frontocentral ERP amplitudes as a function of trial duration for different walking speeds (slow, medium and fast, vertical dashed line indicates stimulus onset). The black bands indicate the time points where the amplitudes are statistically significant (monte carlo cluster corrected *p* < 0.05). Topographies illustrate scalp distribution of EEG activity at the stimulus duration. (B) The linear relationship between PSE and occipital and (E) frontocentral ERPs for different walking speeds.(C) The linear relationship between confidence and onset locked occipital and (F) frontocentral ERPs. Scatterplots depict individual data across different stimulus durations. Shaded gray areas depict standard error (SE). Asterisk indicate significance level (**= *p* < 0.01; *** = *p* < 0.001).

To further disentangle the effect of walking speed, we examined spectral power at occipital and frontocentral electrodes between 0.5-5Hz. Here, normalized spectral power at occipital electrodes revealed three significant clusters (all *p* < 0.05), located at each of the three walking speeds. Planned pairwise comparisons yielded maximum log power at the actual stimulus frequency for each walking speed (all *p* bonferroni< 0.05, except *M*_fast 3 hz - medium 3 hz,_ *p*_bonferroni_ = 0.198 and *M*_medium 1.6 hz - slow 1.6 hz,_ *p*_bonferroni_ = 0.563), indicating that this direct scaling was a result of neural entrainment (Figure 5A & B). Critically, at frontocentral electrodes, the power spectrum significantly differed only for the slow motion frequency, with highest power observed for slow motion (monte carlo cluster corrected *p* < 0.05; Figure 5C & D). Thus, the neural entrainment was only partially observed in the frontocentral electrodes.

**Figure 5.**
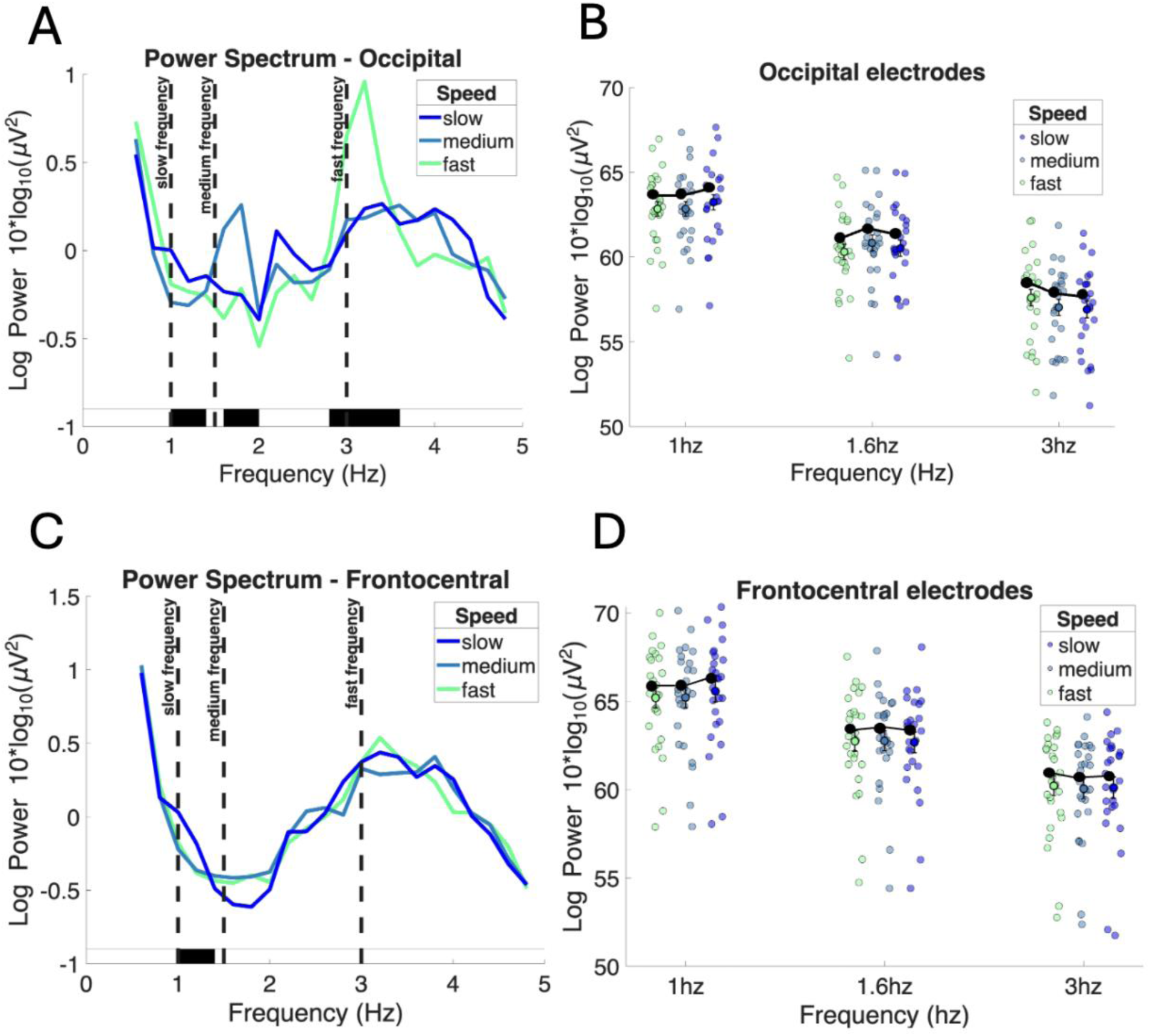
Neural entrainment effect for time dilation across occipital and frontocentral regions. (A & B) Power spectrum for each walking speed for occipital and (C & D) frontocentral electrodes. Error bars indicate standard error (SE).

Further, we investigated the linear relationship between each PSE and onset locked ERPs for each stimulus speed separately for frontocentral and occipital electrodes with the following linear mixed effects model, which included PSE as the fixed slope and participants as random intercept:

Model 6. ERP ∼ PSE + 1|participant

Linear mixed effects model revealed a significant negative relationship between onset locked ERPs from occipital electrodes and PSE values (ß_occipital_ =-0.701, *SE* = 0.17, 95% CI = [− 1.04 −0.36], *p* < 0.001, Figure 4B), indicating a decrease in ERP amplitudes from occipital electrodes as a function of PSE. However, this relationship was not statistically significant for frontocentral electrodes (ß_frontocentral_ = 0.47, *SE* = 0.31, 95% CI = [−0.14 1.082], *p* = 0.131, Figure 4E), with constant ERP amplitudes across different PSEs. Together, these results suggest that the leftward shift in psychometric functions are driven uniquely by occipital regions.

We further investigated the relationship between the onset locked ERPs and confidence ratings associated with each stimulus speed, separately for frontocentral and occipital electrodes with the following linear mixed effects model:

Model 7. ERP ∼ Confidence + 1|participant

Linear mixed effects model revealed a significant negative relationship between onset locked ERPs from frontocentral electrodes and confidence ratings (ß_frontocentral_ = −3.63, *SE* = 1.31, 95% CI = [−6.24 −1.02], *p* = 0.007, Figure 4F), indicating decrease in ERP amplitudes as a function of subjective confidence. On the other hand, this relationship was not statistically significant for occipital electrodes (ß_occipital_ = −0.28, *SE* = 0.83, 95% CI = [−1.93 1.38], *p* = 0.741, Figure 4C). In contrast to the occipital findings with time dilation, these results suggest that the confidence judgments are encoded uniquely by the frontocentral regions.

## Discussion

Our behavioral results replicated the previous finding that increasing walking speeds for a visual stimulus shifted psychometric curves to the left,indicating a time dilation effect (e.g., Karsilar, Kisa & Balci, 2019; Oztel & Balci, 2021). Further, the point of equal confidence, along with the point of equal RTs showed a similar shift as a function of PSE values. These results further validates that the observed temporal dilation effect emerged as a function of increased internal clock speed (and thus, increased sensory uncertainty), rather than being a result of mere response bias (e.g., Gallagher, Suddendorf & Arnold, 2019; Morgan et al., 2012). This idea is also in line with Akdogan and Balci’s (2017) original model predictions such that the two internal clocks that are being compared for metacognitive monitoring should be affected by the stimulus induced changes in the clock speed in the same way, which thus result in metacognitive blindness for stimulus induced temporal illusions (for a similar discussion see: Oztel & Balci, 2020).

Our EEG findings revealed that the onset locked ERP amplitudes from the posterior sites associated with each stimulus duration (i.e., 1-3.5 seconds) peaked around the offset of stimulus duration (Figure 5, similar to Ofir & Landau, 2022; Noguchi & Kakigi, 2006). This ERP pattern that tracks the onset and offset of the timing stimulus suggests that representations associated with sensory time are encoded at the posterior sites of the brain, where visual sensory information is being processed. Furthermore, the occipital ERP amplitudes that were directly scaled with motion velocity were negatively correlated with the shifts in the psychometric functions. However, this linear relationship was not observed with the confidence ratings. Critically, the peaks in the power spectrum at the stimulus frequencies suggest that the time dilation effect that we observed in the current study was a result of neural entrainment of steady-state motion visual evoked potential (SSMVEP; e.g., Yan et al., 2017). This idea complements the previous studies with different experimental approaches using flickering visual stimuli (e.g., temporal reproduction: Hashimoto & Yotsumoto, 2018; temporal bisection: Herbst et al., 2014). Together, these results suggest that the stimulus induced time dilation effect that we observed in the current study (see also: Karsilar, Kisa and Balci, 2019; Oztel & Balci, 2020) emerges via bottom up sensory processes associated with the to-be timed stimulus.

The ERP signals from the frontocentral region including CNV and LPCt amplitudes portrayed a fully distinct pattern compared to the occipital region. First, contrary to our expectations, the frontocentral ERP amplitudes did not show any direct scaling with motion velocity, nor were correlated with the shift in the psychometric functions. Similar results were observed by Herbst and colleagues (2014), where the flicker induced steady-state visual evoked potential (SSVEP) at the occipital region did not translate in any such modulation of CNV amplitudes. Thus, the authors concluded that the time dilation effect induced by visual stimulus properties (i.e., flicker versus constant stimulus) was not due to changes imposed in the encoding processes of stimulus durations, but could rather be associated with changes in memory or decision processes. However, that study only examined frontocentral responses, and so may have missed sensory-evoked ones from posterior electrodes that could have correlated with the effect, as observed here. Thus, complementing Herbst and colleagues’ finding (2014) our results suggest that the time dilation effect is a result of sensory evoked readings that take place uniquely in the occipital regions.

Second, the occipital and frontocentral regions differ in the nature of temporal information they encode: instead of marking stimulus onset and offset as in occipital activity, for longer durations our results exhibited a ramping up of frontocentral activity until around the median stimulus duration and stabilized (similar to: Ng et al., 2011), which then started degrading before the actual stimulus offset, presumably because a decision has been made prior to stimulus offset (for a similar discussion see: Lindberg & Kieffaber, 2013; Wiener & Thompson, 2015; Ofir & Landau, 2022). Critically, this discrepancy between the first degradation time of frontocentral activity and actual stimulus offset prolonged as a function of stimulus duration. While this pattern is a clear demonstration of how subjective time perception is neurally represented, it could be indicative of changes in (1) mere decision processes associated with different durations (as suggested by Herbst et al., 2014; Kononowicz et al., 2016; van Rijn et al., 2011) or (2) changes in internal clock speed (e.g., Penton-Voak et al., 1996). One important note to entangle these two possibilities is that we observed similar behavioral shifts in confidence ratings that followed motion velocity in the same direction as imposed in shifts in psychometric functions. Here, the rationale is that, if the time dilation effect would be imposed by decision bias rather than any representational changes in temporal percepts imposed by different motion velocities, confidence ratings would remain resilient to changes in motion velocity (also see: Gallagher et al., 2019). Ultimately, the same directional shift in confidence as in psychometric functions that indicate time dilation serves as a further validation instrument that dilated temporal percepts can not emerge simply as a result of change in decision parameters.

Third, unlike the occipital ERPs, the frontocentral ERP amplitudes were negatively correlated with confidence ratings, suggesting that the metacognitive readings associated with time dilation is uniquely represented in the frontocentral region which encompasses both SMA (as implicated in subjective time perception: e.g., Coull et al., 2015) and ACC (as implicated in conflict monitoring and metacognitive processing: e.g., Botvinick et al., 2004). Thus, this result highlights the neural dissociation between the sensory time and the metacognitive representations, as well as the bottom up initiation of time dilation at the occipital region. Similar to this, Weaver and colleagues (2019) observed an occipital contribution unique to visual/perceptual decision but that coupled occipital and frontocentral contributions were observed for confidence ratings for these decisions. Complementing this, several other research documented similar frontocentral contribution to metacognitive processes for low level perceptual decisions (e.g., Zakrzewski et al., 2019; CNV predicts confidence: Boldt et al., 2019; central midline activation: Feuerriegel et al, 2022; blood oxygen level dependent (BOLD) activity in right SMA: Gherman & Philiastides, 2015 and ACC: Chen et al., 2013; Fleck et al., 2006; causal involvement of ACC: Stolyarova et al., 2019). Our findings provide further support for the frontocentral contribution to metacognitive readout might have emerged as a top-down sampling of motor output associated with bottom-up sensory information regarding motion velocity.

Our results suggest that the metacognitive inability to track dilated subjective time perception induced via motion velocity is a result of double dissociation between how time dilation and their metacognitive readings are encoded in the brain. This double dissociation can be explained with a dual stream model where the first stream includes posterior/visual regions that encode sensory time associated with the presentation of a visual stimulus. These sensory signals that are exposed to task-irrelevant visual information (i.e., walking speed) could be incorporated in the frontoncentral zone. These visually corrupted temporal representations are integrated in the SMA in a way that eventually entails dilation in the subjective representations of stimulus duration. In the second stream, the dilated time signals might be encoded as conflicting information between the sensory and subjective representations associated with stimulus duration in ACC. On the other hand, given that the sensory representations associated with physical stimulus duration are not readily accessible in ACC, the accurate metacognitive readings for time dilation effect is not possible. This dual stream model can capture how accurate metacognitive evaluation of the stimulus induced time dilation effects is not plausible.

One caveat here is that the dual stream model that we propose in the current study serves as a post hoc explanation of the observed results, rather than being a priori predicted and tested. This includes the involvement of ACC and SMA during metacognitive processing of time dilation effect such that this interpretation is primarily led by the previous studies given that the spatial resolution of EEG is very limited. Thus, future research should devise an experimental approach that directly tests the validity of this explanation and pinpoint exact brain regions that are involved during metacognitive processes associated with time dilation effect to verify the proposed connection between SMA and ACC with brain imaging methods where spatial resolution is enhanced substantially.

In conclusion, the current study replicated the previous behavioral findings that while human time perception is subjected to dilation effects induced by motion velocity associated with timed stimulus, the metacognitive system fails to capture these contextual changes in the subjective temporal representations. This metacognitive inability results in a double dissociation between how time dilation and metacognitive readings associated with them are encoded in the neural level, such that time dilation emerges as a bottom-up sensory influences and metacognition reads out dilated representations via top-down processing of motor output.

## Acknowledgement

This study was supported by the National Science Foundation (NSF) awarded to M.W. (grant number: #2342812)

## Data Availability Statement

Data will be available upon request.

## Use of AI Declaration

Authors confirm that no section of this manuscript has been drafted or rewritten by generative AI tools.

